# Methamphetamine learning induces persistent nonmuscle myosin II-dependent spine motility in the basolateral amygdala

**DOI:** 10.1101/605394

**Authors:** Erica J. Young, Hua Lin, Theodore M. Kamenecka, Gavin Rumbaugh, Courtney A. Miller

## Abstract

Nonmuscle myosin II inhibition (NMIIi) in the basolateral amygdala (BLA) selectively disrupts memories associated with methamphetamine (METH) days after learning, without retrieval. However, the molecular mechanisms underlying this selective vulnerability remain poorly understood. A known function of NMII is to transiently activate dendritic spine actin dynamics with learning. Therefore, we hypothesized that METH-associated learning perpetuates NMII-driven actin dynamics in dendritic spines, leading to an extended window of vulnerability for memory disruption. Two-photon imaging of actin-mediated spine motility in neurons from memory-related structures, BLA and CA1, revealed a persistent increase in spine motility after METH-associated learning that was restricted to BLA neurons. METH-induced changes to BLA spine dynamics were reversed by a single systemic injection of an NMII inhibitor. Thus, a perpetual form of NMII-driven spine actin dynamics in BLA neurons may contribute to the unique susceptibility of METH-associated memories.

---

Dendritic spines are small, actin-rich structures lining dendrites of excitatory neurons. These postsynaptic compartments are dynamic and enable input-specific biochemical and electrical isolation of synapses to facilitate signal transduction and information storage (1–3). During learning, dendritic spines undergo both structural and functional changes to stabilize synapses and, ultimately, memory (4, 5). Consistent with this, there is a tight connection between the physical geometry of spines and the ability to transform experiences into long-term memory (6–8). Polymerization of actin, the elongation and complex branching of filamentous actin (F-actin), drives the spine structural plasticity that is required for functional plasticity and learning (9–14).

Interestingly, long-term potentiation (LTP) and newly formed memories become impervious to actin depolymerization shortly after the underlying synaptic plasticity occurs (15–17). This is attributed to rapid stabilization of the actin cytoskeleton through the cessation of polymerization and recruitment of actin capping and stabilizing proteins (18). However, we recently made the unexpected discovery that memories associated with the commonly abused stimulant, methamphetamine (METH), remain uniquely susceptible to actin depolymerization many days after learning (19). Indeed, a single infusion of the actin depolymerizer, Latrunculin A (LatA) into the basolateral amygdala (BLA) results in an immediate, long-lasting and retrieval-independent loss of the METH-associated memory and associated drug seeking behavior. Furthermore, this memory loss is accompanied by a return of BLA spine density to pre-METH conditioning levels. LatA works by sequestering actin monomers, removing them from the pool available for addition to F-actin during polymerization (20, 21). In this way, LatA influences populations of dynamic, but not stable, actin. Thus the susceptibility of a METH-associated memory days after learning suggests that METH may interfere with the normal actin stabilization mechanisms in BLA spines.

Memories associated with drugs of abuse can elicit drug seeking behavior, even after behavioral modification therapy and prolonged periods of abstinence. This powerful motivation is triggered, in large part, by activation of the amgydala (22–24). Therefore, the ability to rapidly and selectively disrupt amygdala-dependent memories associated with METH represents a potential therapeutic avenue (25). Because actin’s critical role in cellular processes outside the CNS limits its potential as a drug target, we shifted focus to the molecular motor ATPase, nonmuscle myosin II (NMII), a driver of actin polymerization (26, 27). NMII is a hexameric protein made of two heavy chains (MHC), two regulatory light chains and two essential light chains. The MHCs are the workhorse of NMII, bearing both the ATPase and actin-binding sites, with the motor heads binding to actin and moving it through physical force. We have previously demonstrated that NMII bearing the IIB MHC (*Myh10*) is a major driver of synaptic actin polymerization to promote and stabilize LTP and long-term memory formation in CA1 of the dorsal hippocampus and the BLA (15, 28). In addition, through genetic and pharmacologic manipulations we have established that targeting NMII either directly in the BLA or systemically is well-tolerated and recapitulates the METH-associated memory, drug seeking and BLA spine density effects of direct actin depolymerization (29, 30).

The rapid and persistent impact of NMII inhibition on METH-associated memory is surprisingly specific, having no such effect on memories associated with foot shock, food reward, or other drugs of abuse (nicotine, morphine, cocaine or mephedrone) (31). The underlying mechanism responsible for this selectivity is largely unknown to date. The impact of actin depolymerization on BLA spine density via LatA or NMII inhibition with Blebbistatin (Blebb) indicates a connection to spine actin. Therefore, we hypothesize that METH-associated memory’s selective susceptibility is due to NMII-dependent sustained actin dynamics in spines. However, actin is present in many compartments of the cell, including the presynapse and nucleus. Therefore, as a first step towards addressing our hypothesis that BLA spine actin is persistently altered by METH conditioning, we utilized time-lapse, two-photon imaging of BLA spine motility, the extension and retraction of the spine head from the dendrite. These actin-dependent structural dynamics have been recorded in a number of brain regions, including cortex, hippocampus and cerebellum, both in cultured and acute slices, and in vivo during early development and in response to synaptic stimulation (32–39). Actin-driven spine motility is also age-dependent, with basal motility rates highest in early life, as it likely acts as an important component of synaptic wiring during development (32). In support of this, disruption of critical periods by visual deprivation or via genetically linked neurodevelopmental disorders, such as *SYNGAP1* mutation, results in premature decreased spine motility and profound effects on adult cognitive function (36, 40). Additional evidence indicates that motility related to the spine neck specifically regulates spine calcium decay kinetics, thus dynamically influencing the plasticity potential of individual spines (41, 42).

Here we assessed spontaneous spine motility in the BLA for the first time, as well as motility after METH conditioning and the impact of NMII inhibition. We also assessed CA1 spine motility for comparison. Like the BLA, the dorsal HPC is a critical component of the neural circuit supporting drug-associated memories and we have reported spine density increases in CA1 with METH learning, as in the BLA (31, 43). However, METH memories do not bear the same vulnerability to disruption when NMII is specifically inhibited in CA1 and, unlike BLA spines, METH-induced CA1 spine density increases remain intact after systemic Blebb treatment (31). Therefore, we further hypothesized that METH conditioning would result in an NMII-dependent increase in spine motility in the BLA, but not CA1. Indeed, we report that BLA spontaneous spine motility is similar to other brain regions in terms of its actin-dependence and decreased rate with development. Further, METH conditioning increased motility in BLA, but not CA1 spines and this was reversed when animals were treated with systemic Blebb prior to imaging.

## RESULTS

### BLA and CA1 spine motility is actin-dependent and decreases with development

Spine motility has not been assessed in the BLA. Therefore, we first examined age-dependent spine motility, as it has been reported to decrease with development elsewhere in the brain. The exact age at which spine motility decreases varies by brain region and preparation (34, 36, 39, 40). Therefore, we examined spine movements across two postnatal (P) ranges, days P16-21 and P28-35. Acute slices from naïve Thy1-GFP(m) mice, containing both CA1 and BLA, were imaged every 5 min for one hour. The motility of 30 spines per slice was then quantified. Consistent with prior reports on spine motility in the hippocampus (35, 39), CA1 spines displayed more motility at P16-21, as compared to P28-35 (**Fig. 1A**: T_(12)_ = 3.77, *P* < 0.01; Power = 0.93). Prior to performing any additional analyses, we confirmed that the range of spine motilities observed as a group was represented across spines from all slices analyzed, and was, therefore, not biased by a slice with particularly high or low overall motility (**Fig. 1B**). This was performed for all experiments in the study. To determine whether the shift in spine movements was driven by decreased motility across the whole population of spines or by loss of a subgroup of spines only present in P16-21 CA1, we examined the cumulative distribution of spine movement. This revealed a shift in the whole population of spines towards decreased movement at P28-35, as well as the loss of highly motile spines (movement ≥0.030 μm/min; **Fig. 1C**; KS Test *P* < 0.0001). BLA spine movements followed a similar developmental pattern (**Fig. 1D**; T_(14)_ = 5.01, *P* < 0.001, Power = 0.99), with an even greater shift in the population of spines towards decreased motility at P28-35 (**Fig. 1F**, KS test *P* < 0.0001).

**Figure 1.**
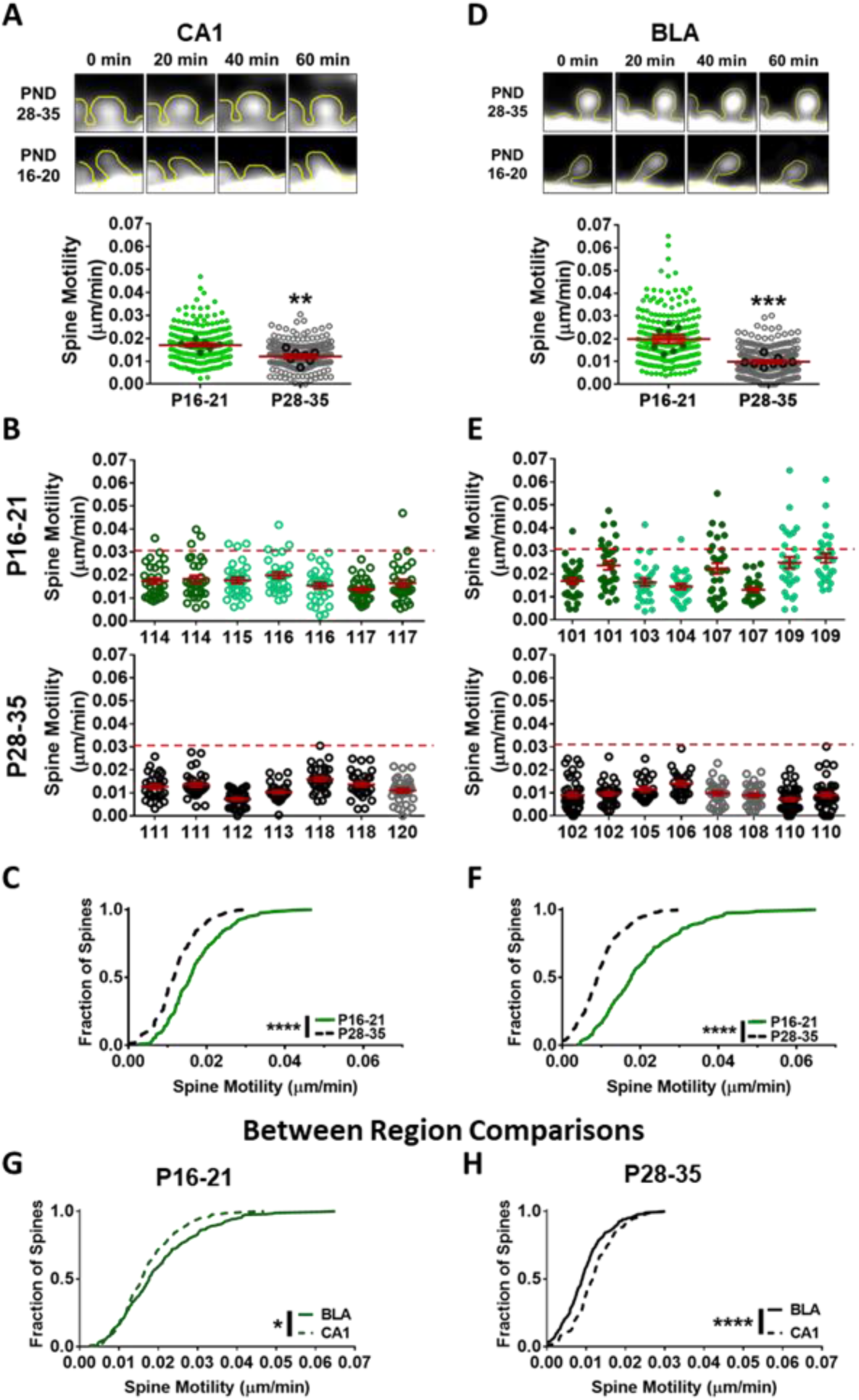
Motility of CA1 and BLA spines decreases with age. (A) CA1 dendritic spine motility from naïve young animals (P16-21) and early adolescents (P28-35). Small lighter colored circles represent each individual spine’s movement over one hour, while the large darker circles represent the average of 30 spines from each slice. (P28-35 *n* = 210 spines, 7 slices, 5 animals; P6-21 *n* = 210 spines, 7 slices, 4 animals) (B) CA1 spine motility organized by animal and slice. Slices from female animals are in light green and grey, males are depicted by dark green and black. (C) Cumulative distributions of CA1 spine movements from P16-21 and P28-35. (D-E) Spine movements measured in BLA slices from P16-21 and P28-35, and (F) corresponding cumulative distributions. (P28-35 *n* = 260 spines, 8 slices, 5 animals; P16-21 *n* = 240 spines, 8 slices, 5 animals) (G-H) Comparison of BLA and CA1 spine movements at P16-21 and P28-35. * *P*<0.05 and **** *P*<0.0001.

We next performed unbiased cluster analysis, identifying three clusters. This enabled comparison of the relative representation of spines in low-, mid- and high-range motility within CA1 and BLA in early life (P16-21) versus early adolescence (P28-35; combined overall χ^2^_(6)_ = 95.93, *P* < 0.0001; CA1 overall χ^2^_(2)_ = 20.30, *P* < 0.0001; BLA overall χ^2^_(2)_ = 65.82, *P* < 0.0001 **Fig. S1A**). Chi-squared analysis established age-dependent differences in CA1 for low (Cluster 1) and mid-range (Cluster 2) motility (Cluster 1 χ^2^_(1)_ = 20.15, *P* < 0.0001; Cluster 2 χ^2^_(1)_ = 19.14, *P* < 0.0001; Cluster 3 χ^2^_(1)_ = 1.00, *P* > 0.05 **Fig. S1B**), while motility decreased in all three clusters (including high motility) with age for BLA spines (Cluster 1 χ^2^_(1)_ = 65.64, *P* < 0.0001; Cluster 2 χ^2^_(1)_ = 57.43, *P* < 0.0001; Cluster 3 χ^2^_(1)_ = 6.58, *P* < 0.01; **Fig. S1C**). When comparing regions, BLA spines were more motile than CA1 spines during early life (P16-21; KS *P* < 0.05), while the opposite was true in early adolescence (P28-35; KS *P* < 0.0001; **Fig. 1G-H**).

Additional similarities and differences were observed between CA1 and BLA. The most pronounced difference was in initial spine lengths. BLA spines were longer than those in CA1 and this was not age-dependent (**Table 1**). This is consistent with a prior report using Golgi staining, though a direct comparison was not made (44). Examining individual spines in early development (P16-21), a time point when motility was sufficiently elevated to reveal variability in movements between spines over the course of an hour, we found CA1 and BLA spines with low to medium overall motility displayed a remarkably similar range of movement events (**Fig. 2A-B**). These movements were relatively infrequent and represented small to medium changes in length, rather than, for example, only 1-2 large movements that averaged to moderate motility over the course of an hour. Differences emerged in spines with overall high motility rates; BLA spines made larger, more frequent movements when compared to CA1 spines (top panel). In addition, the appearance and disappearance of spines over the course of imaging, while rare, was observed in both regions. It was more prominent in young BLA slices and CA1 slices from both age ranges, occurring at 3% of spines.

**Figure 2.**
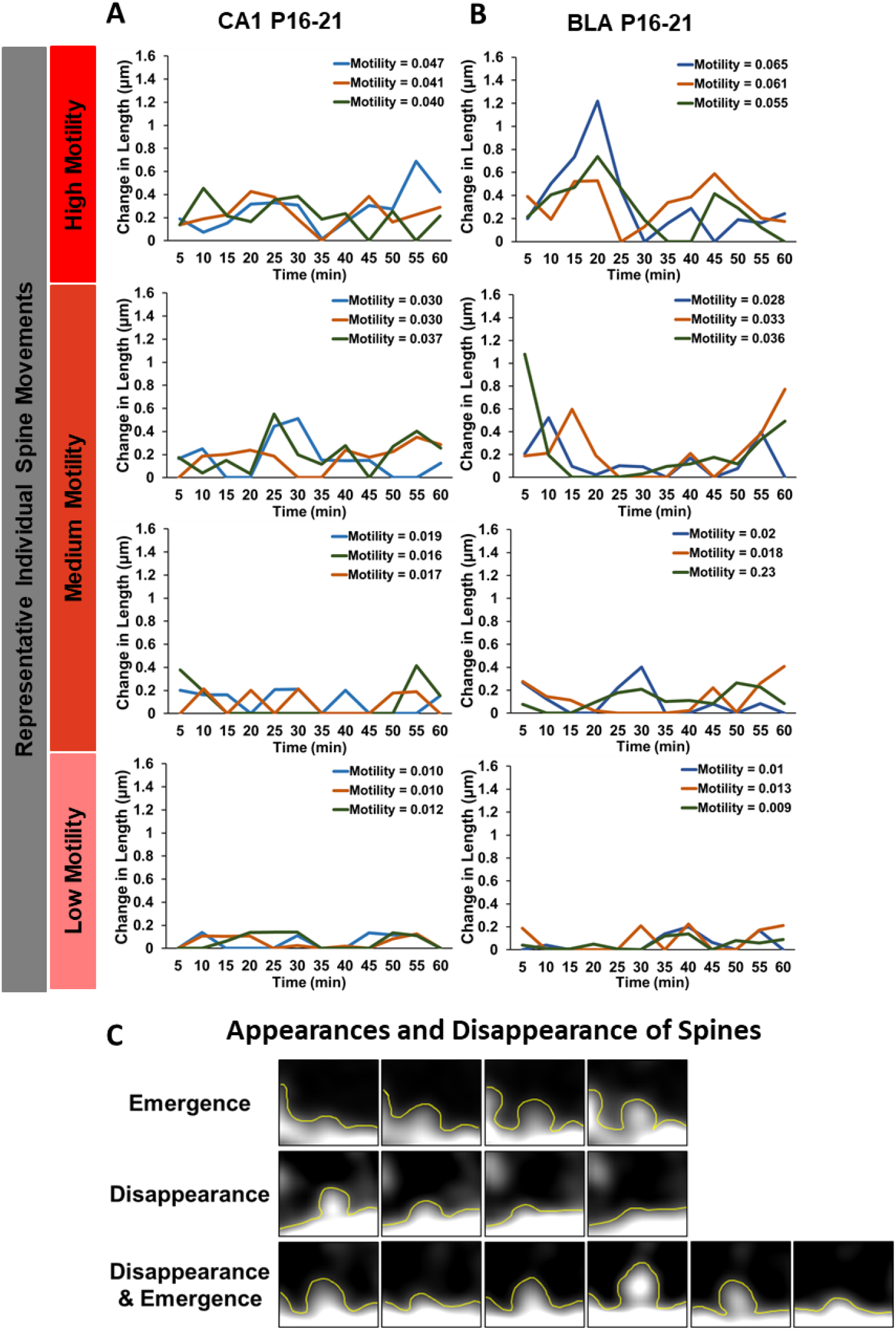
Age-related CA1 and BLA spine movements. Representative movements of individual (A) CA1 and (B) BLA spines with different levels of motility over one hour. (C) Examples of spines emerging and disappearing during the imaging period.

**Table 1.**

| Initial Spine Length |  |  |  |  |
| --- | --- | --- | --- | --- |
| Age-Related Comparisons<br>(within regions) |  |  |  |  |
|  | BLA | Statistics<br>Mann-Whitney | CA1 | Statistics<br>Mann-Whitney |
| P28-35 | 1.44 ± 0.03 | U = 29522<br>P = 0.299 | 1.30 ± 0.02 | U = 21603<br>P = 0.719 |
| P16-21 | 1.52 ± 0.03 |  | 1.30 ± 0.02 |  |
| Region-Related Comparisons<br>(between regions) |  |  |  |  |
|  | Region | Mean |  | Statistics<br>Mann-Whitney |
| Young<br>(P16-21) | BLA | 1.52 ± 0.03 |  | U = 19580<br>P < 0.0001 |
|  | CA1 | 1.30 ± 0.02 |  |  |
| Adolescent<br>(P28-35) | BLA | 1.44 ± 0.03 |  | U = 22676<br>P = 0.0016 |
|  | CA1 | 1.30 ± 0.02 |  |  |

Spine motility is actin-dependent in the hippocampus and cortex (45, 46). Therefore, we next determined the actin dependence of BLA spine dynamics via bath application of the actin stabilizer, jasplakinolide (Jasp), to P16-21 slices. Jasp decreased BLA spine movement (**Fig. 3A-B**, T_(12)_ = 5.48, *P* = 0.0001, Power = 0.99; KS test *P* < 0.0001). This included an absence of highly motile spines, making it more akin to P28-35 spine motility distribution (**Fig. 2F**). We also confirmed there were no group differences in initial spine length for this and all subsequent experiments (**Table 1 and Table S1**). Taken together, these data establish that BLA spine motility is actin-dependent and decreases markedly by P28-35, with a significant loss of highly motile spines (≥0.03 μm/min).

**Figure 3.**
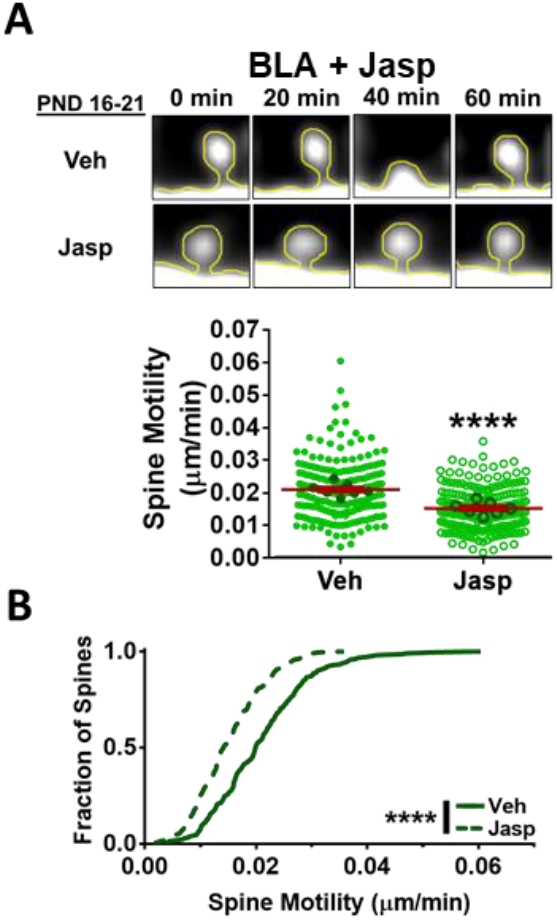
Actin Stabilization of young BLA spines. Young (P16-21) BLA slices were incubated in bath applied Veh or Jasp for 30min prior to imaging. Spine movements and cumulative distributions were measured in the presence of Jasp or vehicle. (Veh *n* = 210 spines, 7 slices, 5 animals; Jasp *n* = 210 spines, 7 slices, 6 animals). Error bars represent SEM and ** *P*<0.01, *** *P*<0.001, **** *P*<0.0001.

### METH conditioning increases basal spine motility in BLA, but not CA1

We next assessed BLA spine motility following METH-associated learning as a measure of spine actin dynamics. We have previously demonstrated that METH-associated memory is vulnerable to disruption by Blebb days after training in adult and adolescent (P28-35) male and female mice (30). In the current study, P28-35 male and female Thy1-GFP(m) mice underwent conditioned place preference (CPP) training with either saline or METH, followed by imaging one to three days later (**Fig. 4A**). Similar to P16-21 motility in naïve slices (**Fig. 2**), differences in individual spine movements were the most pronounced as motility increased (**Fig. S2 A-B**). Additionally, as in naïve slices, the appearance and disappearance of spines over the course of imaging was rare following CPP training (1.7-2.9%) and was not influenced by METH exposure.

**Figure 4.**
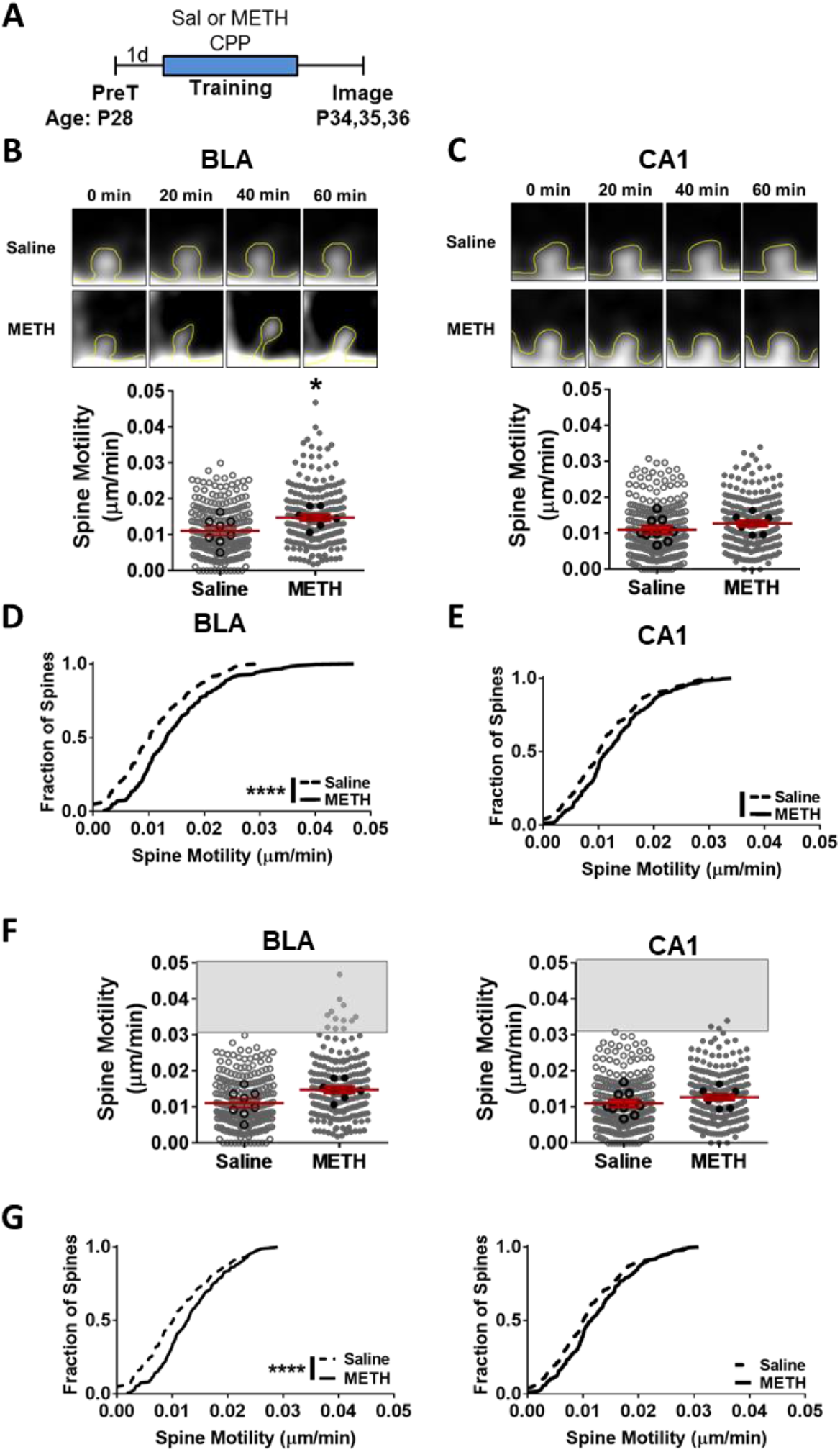
METH conditioning produces a persistent increase in spine motility in BLA, but not CA1. (A) Schematic of experimental design. Spine movements in the BLA (B) and CA1 (C) were measured over one hour, 1 to 3 days after Saline or METH training. Small circles represent each spine’s movement, while the larger circles with heavier borders are the average of 30 spines per slice. (BLA: Saline *n* = 240 spines, 8 slices, 8 animals; METH *n* = 240 spines, 8 slices, 7 animals; CA1: Saline *n* = 270 spines, 9 slices, 7 animals; METH *n* = 240 spines, 8 slices, 6 animals) Cumulative distributions of spine movements in the BLA (D) and CA1 (E) following training. (F) ROC curves were used to determine cut off for highly motile spines (those in the grey boxes). The cut-off was set as the maximum motility displayed by any spine from the Saline-treated condition (0.030 μm/min). (G) Cumulative distributions of spine movements with highly motile spines removed. Error bars represent SEM, ** *P*<0.01, *** *P*<0.001, **** *P*<0.0001.

Spine motility was elevated in the BLA, but not CA1, 1-3 days after METH training relative to saline controls (**Fig. 4B-C**; BLA T_(15)_ = 2.38, *P* < 0.05, Power = 0.60; CA1 T_(16)_ = 1.27, *P* > 0.05, Power = 0.22) and motility was equally represented across slices analyzed (**Fig. S2C-D**). Moreover, every slice imaged from METH trained animals, had spines in the BLA that reached the “high motility” threshold of ≥ 0.03 μm/min (**Fig. S2C**), identified in Figure 1’s age-dependent motility experiment. As a reminder, the BLA and CA1 of young (P16-21), but not adolescent (P28-35) animals displayed spine motility rates above 0.03 μm/min (**Fig. 1D-F**). High motility spines were not observed in the BLA from the saline condition (**Fig. S2C**) or in CA1 following saline or METH training (**Fig. S2D**). Interestingly, METH-related increases in BLA spine motility did not correspond to increases in spine width, when assessed in a subset of spines with high and average motility (T_(27)_ = 0.92, *P* > 0.05; data not shown).

Further analysis of spine motility with a KS test of the cumulative distribution of spines revealed that METH training shifted all spine movements in the BLA (*P* < 0.0001), but not CA1 (*P* = 0.06), toward higher motility and precipitated the appearance of highly motile spines (**Fig. 4D-E**). Even with highly motile spines identified and removed using ROC to establish a cut-off criteria, METH training still significantly increased BLA spine movements (BLA *P* < 0.0001, Power = 1.0; CA1 *P* > 0.05, Power = 0.82; **Fig. 4F-G**). Thus the effect of METH on BLA spine dynamics cannot be attributed solely to a subset of highly dynamic spines. However, given the sparse distribution of memories it is plausible that METH-associated memory is preferentially targeted to highly motile spines.

To examine motility in greater depth, cluster analysis was performed (Combined overall χ^2^_(6)_ = 33.29, *P* < 0.0001; BLA overall χ^2^_(2)_ = 15.07, *P* < 0.001; CA1 overall χ^2^_(2)_ = 6.58, *P* < 0.05 **Fig. 5A**), as in Fig. S1A. Chi-squared analysis showed that METH decreased the number of spines in the low motility group (Cluster 1 χ^2^_(1)_ = 9.25, *P* < 0.01) and increased the number of spines in the high motility group (Cluster 3 χ^2^_(1)_ = 10.59, *P* < 0.001), relative to spines from saline-treated mice (Cluster 2 χ^2^_(1)_ = 3.33, *P* > 0.05 **Fig. 5B**). In contrast, CA1’s high motility Cluster 3 was unchanged by METH training (Cluster 1 χ^2^_(1)_ = 5.59, *P* < 0.05; Cluster 2 χ^2^_(1)_ = 4.11, *P* < 0.05; Cluster 3 χ^2^_(1)_ = 2.20, *P* > 0.05 **Fig. 5C**). Interestingly, BLA spine movement appeared to be restricted to the spine head, as very little neck motility was observed in spines, regardless of treatment during training, and it did not correlate with the overall motility rate of the spine (F_(3, 116)_ = 20.61, *P* < 0.0001; Neck Saline vs METH *P* > 0.05; Overall length Saline vs METH *P* < 0.01; **Fig. S3**). Collectively, these data indicate that METH-associated learning produces a selective and lasting increase in the basal dynamics (no acute stimulation at the time of imaging) of P28-35 BLA spine heads that are normally quite stable.

**Figure 5.**
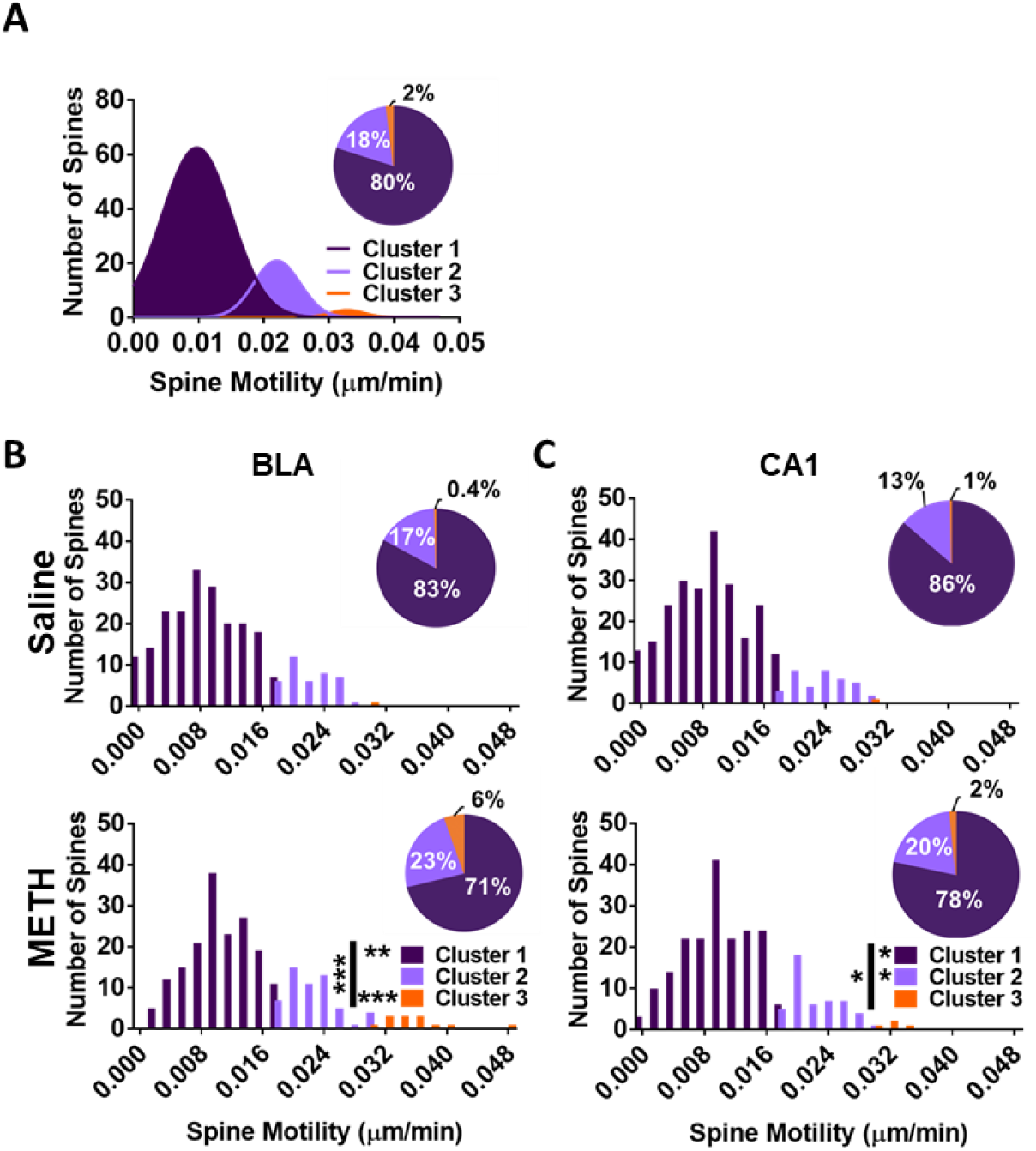
Cluster analysis of spine motility following CPP training. (A) All BLA and CA1 spines underwent cluster analysis. Number of spines in each cluster for (B) BLA and (C) CA1 spines. Error bars represent SEM, ** *P*<0.01, *** *P*<0.001, **** *P*<0.0001.

### Persistent METH-induced BLA spine motility is NMII-dependent

Systemic inhibition of NMII by Blebb disrupts METH-associated memory and returns BLA spine density to pre-METH, baseline levels. Therefore, we next determined the effect of Blebb on BLA and CA1 spine motility to test the hypothesis that METH-induced BLA spine dynamics are sustained by NMII. Animals underwent saline or METH-associated CPP training, as in Figure 4. One to three days later and 24 hours prior to imaging, mice were treated with either Blebb or vehicle (IP), resulting in four groups (Saline/Veh, Saline/Blebb, METH/Veh, METH/Blebb; **Fig. 6A**). Comparing the spine motility of saline and METH-trained animals given vehicle 1-3 days post-training replicated the results reported in Figure 2. Specifically, METH conditioning increased the overall motility of spines in the BLA, but not CA1 (BLA F_(3, 25)_ = 6.69, *P* < 0.01, METH/Veh vs Saline/Veh, Saline/Blebb, METH/Blebb *P* < 0.001, Power = 0.95; CA1 F_(3, 24)_ = 0.95, *P* > 0.05, Power = 0.23; **Fig. 6B-C**). The cumulative distribution of spines followed a similar pattern, with a large shift in BLA motility and increased representation of highly motile spines (**Fig. 6D**, top right panel; Saline/Veh vs METH/Veh *P* < 0.0001). CA1 spines showed a small, but significant shift in the distribution of motility, but no appearance of highly motile spines (**Fig. 6E**, top right panel; Saline/Veh vs METH/Veh *P* < 0.05). Again, all effects were broadly consistent across slices (**Fig. S4**). Using a protocol that disrupts METH-associated memory and reverts BLA spine density to pre-METH levels (IP Blebb 1-3 days post-training (30)) reversed METH-associated increases in BLA spine movements (**Fig. 6B,D** bottom left panel and **Fig. S4**; METH/Veh vs METH/Blebb *P* < 0.0001). As a result, the distribution of spine motility for the METH/Blebb group was identical to BLA spines from Saline/Blebb group (**Fig. 6D** bottom right panel; Saline/Blebb vs METH/Blebb *P* > 0.05). Blebb had no effect on the motility of BLA or CA1 spines from saline-trained animals (Fig. 6B-C **and** D-E top left panels; BLA: Saline/Veh vs Saline/Blebb *P* > 0.05; CA1: Saline/Veh vs Saline/Blebb *P* > 0.05). While Blebb had no effect on overall spine motility in CA1 (**Fig. 6C**), it did reverse the small rightward shift present in METH-trained, vehicle-treated mice (**Fig. 6E**, bottom left panel; METH/Veh vs METH/Blebb *P* > 0.05). This is further supported by cluster analysis (**Fig. S5**), which revealed that METH training influenced all three Clusters in the BLA, including the development of highly motile Cluster 3 and that this population of spine motility was almost entirely lost with Blebb (**Fig. 6F** and **Fig. S5**; BLA overall χ^2^_(6)_ = 63.68, *P* < 0.0001; Saline/Veh vs Saline/Blebb: Overall χ^2^_(2)_ = 0.01, *P* > 0.05; Saline/Veh vs METH/Veh: Overall χ^2^_(2)_ = 34.63, *P* < 0.0001, Cluster 1 χ^2^_(1)_ = 26.60, *P* < 0.0001, Cluster 2 χ^2^_(1)_ = 13.17, *P* < 0.001, Cluster 3 χ^2^_(1)_ = 16.63, *P* < 0.0001; METH/Veh vs METH/Blebb: Overall χ^2^_(2)_ = 23.25, *P* < 0.0001, Cluster 1 χ^2^_(1)_ = 17.89, *P* < 0.0001, Cluster 2 χ^2^_(1)_ = 8.51, *P* < 0.01, Cluster 3 χ^2^_(1)_ = 11.23, *P* < 0.001; Saline/Blebb vs METH/Blebb χ^2^_(2)_ = 3.32, *P* > 0.05). The most notable difference between the BLA and CA1 following METH training was the complete absence of highly motile spines in CA1 (**Fig. 6F-G** and **Fig. 4F-G**; CA1 Overall χ^2^_(6)_ = 7.24, *P* = 0.06; Saline/Veh vs Saline/Blebb χ^2^_(2)_ = 0.05, *P* > 0.05; Saline/Veh vs METH/Veh χ^2^_(2)_ = 5.34, *P* < 0.05; METH/Veh vs METH/Blebb χ^2^_(2)_ = 3.86, *P* < 0.05; Saline/Blebb va METH/Blebb X^2^_(2)_ = 0.01, *P* > 0.05).

**Figure 6.**
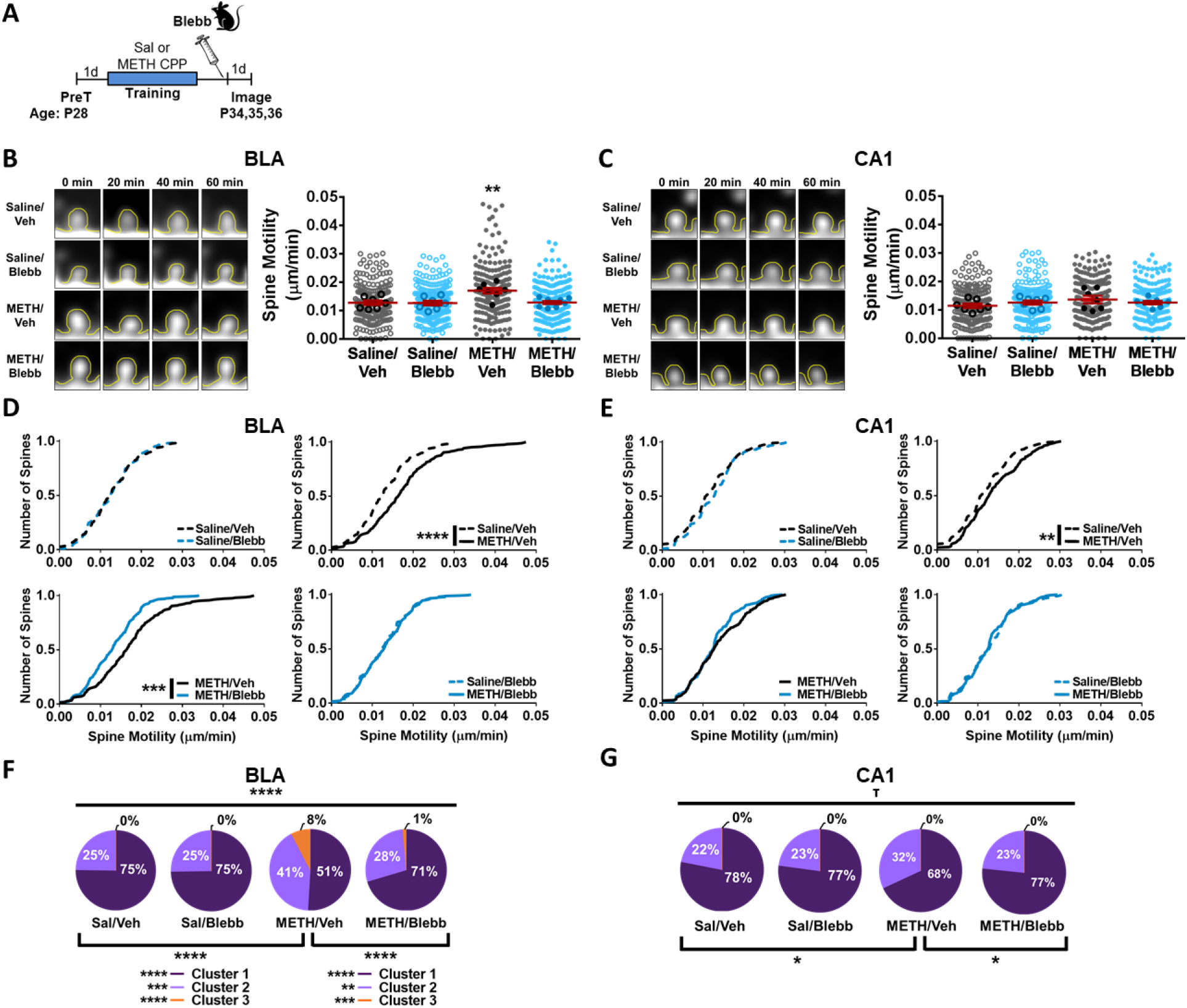
Persistent METH-induced BLA spine motility is NMII-dependent. (A) Schematic of experimental design. (B) Spine movements and representative images of P28-35 BLA slices 24hrs after treatment (Saline/Veh: *n* = 210 spine, 7 slices, 6 animals; Saline/Blebb: *n* = 210 spines, 7 slices, 7 animals; METH/Veh: *n* = 210 spines, 7 slices, 7 animals; METH/Blebb *n* = 240 spines, 8 slices, 7 animals) (C) Representative images and CA1 spine movements (Saline/Veh: *n* = 210 spine, 7 slices, 6 animals; Saline/Blebb: *n* = 210 spines, 7 slices, 5 animals; METH/Veh: *n* = 210 spines, 7 slices, 7 animals; METH/Blebb *n* = 210 spines, 7 slices, 6 animals) Cumulative distribution plots comparing spine motility in the (D) BLA and (E) CA1 of different treatment groups. Summary of cluster analysis of spine movements in the (F) BLA and (G) CA1. Error bars represent SEM and ** *P*<0.01, *** *P*<0.001, **** *P*<0.0001.

## DISCUSSION

Previously, we have reported that direct actin depolymerization or NMII inhibition within the BLA results in an immediate, long-lasting and retrieval-independent disruption of METH-associated memory, as well as a reversal of METH-induced spine density increases to basal levels. Interestingly, NMII inhibition within CA1 does not produce the same retrieval-independent disruption of METH-associated memory, nor does it alter METH-induced increases in spine density. Here we have used two-photon serial, time-lapse imaging of spine motility to begin to address our hypothesis that the selective susceptibility of METH-associated memory and spines to NMII inhibition is due to uniquely persistent, NMII-dependent actin dynamics in BLA spines.

Because this was the first assessment of spine motility in the BLA, we first confirmed that motility rates decrease with age, as has been reported in cultured hippocampal cells and slices, dentate gyrus and CA1 acute slices, and primary visual cortex in vivo (34, 35, 37–40, 45). This also allowed us to compare the rates of spine motility between the BLA and CA1 in early postnatal life and adolescence. BLA spines, which are longer than CA1 spines at both age ranges, were more motile than CA1 at P16-21. This is not overly surprising, as the amygdala undergoes intense innervation between P18-20 (47, 48), a process that occurs earlier in the hippocampus (P14-16; (49)). Therefore, it is possible that more similar spine movements would be observed if P16-21 BLA spines were compared to P14-16 CA1 spines. At P28-35, the pattern reversed, with BLA spines becoming more stable than those in CA1, making the impact of METH training on BLA motility at this age even more striking. In this series of experiments, we also confirmed that spine motility is actin-dependent in the BLA, as has been shown in cultured hippocampal cells and slices (35, 37, 39, 45). This established that spine motility is an appropriate measure to determine the impact of METH on spine actin dynamics.

METH-associated learning produced an increase in the dynamics of P28-35 BLA spines that was present at least three days after the last training session with METH. This is in marked contrast to spines from saline-treated animals, whose BLA spines are quite stable, as well as spines in CA1, regardless of treatment (saline or METH). Interestingly, the spine movements were restricted to the head, as there was no change in motility of the spine neck. Changes in spine neck motility have been connected to functional changes in spines through the regulation of calcium entry and exit (50), suggesting such dynamic changes are not occurring in BLA spines altered by METH training. This is consistent with the notion that the persistent motility in response to METH training does not reflect a process actively involved in the maintenance of the METH-associated memory. Rather, we favor the interpretation that METH-associated memory persists *in spite* of this lack of normal actin-myosin stabilization and that this aberrant plasticity imparts a unique vulnerability of METH-associated memory to disruption long after learning.

NMII inhibition disrupts METH-associated memory and reverses BLA spine density with equal efficacy to direct actin depolymerization via Latrunculin A (19, 29–31). If the persistent BLA spine motility changes observed here are connected to the memory disruption, we would predict that the motility is NMII-dependent. To test this, we METH trained animals, followed by systemic administration of Blebb one to three days later, a time point and manipulation that selectively disrupts METH-associated memory and BLA spine density. Consistent with our hypothesis, NMII inhibition reverted BLA spine motility to control levels, without influencing CA1 motility. In addition, we replicated the effects of METH training on BLA motility reported in Figure 4. It is important to note that Blebb had no effect on BLA spine motility under control (saline-treated) conditions. It is possible that this reflects a dose-dependent effect; that is, a higher concentration of Blebb would disrupt control motility. Higher doses of Blebb are not tolerated by animals, preventing us from directly testing this possibility. However, we are currently developing analogs of Blebb with improved safety profiles that will be used to address this in the future. An alternative explanation to the possibility of a dose-dependent effect is that NMII is specifically involved in the dynamic changes to the spine actin cytoskeleton and the remaining dynamics are driven by actin treadmilling that does not require NMII or by microtubule polymerization (51–53). This would argue for the particular importance of more motile spines. Indeed, approximately 75% of all BLA spines display low motility at P28-35 (<0.015 μm/min) following saline training, with or without Blebb, and the remaining 25% display moderate motility (0.015-0.03 μm/min; **Fig. 3F**). However, METH training shifts the distribution of BLA spines such that moderately and highly motile spines (Clusters 2 and 3; 0.015-0.05 μm/min) account for 50% of the total spines assessed. Remarkably, administering Blebb after METH training results in a very precise return of spine motility distribution to control levels, not below (**Fig. 3D**, bottom right cumulative distribution plot). The notion, that NMII may not contribute to the low level motility seen under control conditions is consistent with prior evidence from our group that NMII is recruited by synaptic stimulation, rather than serving a housekeeping function (15). This is precisely what makes the sustained susceptibility of METH spines to Blebb of interest, as it suggests that NMII remains constitutively active days after METH training. NMII’s ability to drive actin polymerization is dependent upon two critical steps. First, NMII’s regulatory light chain (RLC) must be phosphorylated to drive a conformation change that exposes the head of NMII’s heavy chain (MHC), which contains the ATP binding site (54). ATP provides the energy for the MHC’s motor head to physically slide actin filaments, driving cytoskeletal rearrangement (55). Blebb works by interfering with the ATP binding site (56). However, Blebb can only influence NMII function if the protein is activated via RLC phosphorylation. Thus, Blebb’s ability to disrupt actin-dependent spine motility days after the stimulation events of METH-associated learning suggests NMII’s RLC may need to remain phosphorylated. This could represent a point of divergence between METH-associated memory representation in the BLA versus CA1, as well as METH-associated memory versus memories for other conditioned stimuli. Future studies will be directed at identifying the upstream signaling cascade regulating METH-induced NMII RLC phosphorylation, focusing on the possibility that METH recruits a unique set of players in the BLA that interferes with the normal, post-learning mechanisms responsible for actin-myosin stabilization.

## METHODS

All animal procedures were conducted in accordance with the Scripps Research Animal Care and Use Committee and national regulations and policies.

### Animals

Male and female heterozygous Thy-GFPm mice (Jackson Laboratory) were bred onsite. All animals were weaned at P22-23 and handled three days priors to the start of training on P28.

### Drug

For actin stabilizing imaging experiments, 200nM Jasp (Tocris) in 0.02% DMSO was bath applied to slices. During training, mice received 1mg/kg doses of methamphetamine hydrochloride (Sigma-Aldrich). For systemic Blebb infusions mice received a 10mg/kg (IP) dose of racemic blebbistatin (TSRI) diluted to 1 mg/kg in a vehicle of 0.9%DMSO/25% Hydropropyl β-Cyclodextrin (HPβCD). Vehicle animals received the vehicle without Blebb.

### Behavior

#### Conditioned Place Preference

Adolescent mice were trained as previously described (30). CPP consisted of two phases, pretesting and training, followed by imaging 1 to 3 days later. Pretesting was conducted over 2 consecutive days. Animals received an IP injection of saline before freely exploring all three CPP chambers for 30 min. Either the white or black chamber was assigned as each animal’s METH-paired chamber (conditioned stimulus; CS+) based on their least preferred chamber during the final 15 min of the second pretest session. There were no initial differences between groups for the amount of time spent in either the white or black chamber. Over the next four consecutive days, animals were trained twice daily in 30 min training sessions. Animals either received saline in the CS-chamber in the mornings and METH in the CS+ in the afternoons or were assigned the opposite training schedule. One to three days following the final day of training animals were euthanized for acute tissue collection.

### Spine Motility

#### Acute Slice Preparation

Acute coronal brain slices (350 μm thick) were extracted and sliced in cold cutting solution composed of (mM): 119 Choline Cl, 22 D-Glucose, 4.3 MgSO4, 2.5 KCl, 1 NaH2PO4, 1 CaCl2 and 26.2 NaHCO3. Adolescent animals (PND 28-36) were gravity perfused with cold cutting solution for 40 sec prior to brain extraction and slicing. Following slicing, brain slices were continuously bubbled with 95% O_2_/5% CO_2_ and incubated at 34 °C for 30 min then room temperature for 30 min in artificial cerebrospinal fluid (aCSF) composed of (mM): 119 NaCl, 2.5 KCl, 1.3 MgSO4, 2.5 CaCl2, 1 NaH2PO4, 0.2 Trolox, 11 glucose and 26.2 NaHCO3. For imaging, slices were continuously perfused with oxygenated aCSF at 1.5-2 ml/min.

#### Two-Photon Imaging

Two-photon imaging of the BLC and CA1 was conducted with a multiphoton laser scanning microscopy (Olympus FV1000MPE-TWIN), equipped with a water immersion objective lens (ULTRA 25x, numerical aperture 1.05, Olympus) and Fluoview software (Olympus). To excite eGFP we used a Ti:sapphire laser and collected emitted photons that passed through a 500-550 nm bandpass filter with an external nondescanned detector. To measure spine dynamics, we imaged slices in XYZT dimension (45-60 μm Z stack; 60 min time series; 5 min between stacks) (Clement et al., 2012). The imaging parameters used were excitation = 930nm, power at sample = ~15mW, pixel dwell time = 2 μs, x-y scaling = 0.201 μ/pixel, and z-scale = 1 μm; no averaging. For the effects of actin stabilization, vehicle (aCSF with 0.02% DMSO) or aCSF + Jasp was bath applied to slices. Slices were allowed to stabilize in the imaging chamber and incubate for 30 min prior to imaging. Slices were then imaged in the XYZT dimension over one hour with 5 min in between each 45-60 μm Z stack.

#### Spine Movement Analysis

Spine movements were assessed in both the CA1 and BLC following standard procedure (36, 40). ImageJ software (NIH) was used to process and align images. After Z stack compression of the 12 images (5 min between frames) the resulting two-dimensional projections were aligned to compensate for inherit x-y drifting during time series imaging using the ImageJ plugin, StackReg. Spines used for analysis were selected in an unbiased way with the researcher blinded to the group. Tertiary dendritic segments that were relatively clear of random axonic and dendritic processes were selected for analysis. Along these segments the movement of the first 10 unobstructed dendritic spines was analyzed for a total of 30 spines per slice. The spine length was measured from the base to the tip of the protrusion in each of the 12 images. Spine movement was expressed as the average change in spine length per minute (μm/min) (34, 36, 40).

#### Statistical Analysis

For all studies, the experimenters were blind to the treatment group. The averages of spine movement per slice were found to be normally distributed but all spines per group was not. Therefore, comparisons of groups using slice averages was conducted using T-tests or one-way ANOVAs, while population comparisons were done with Kolmogorov-Smirnov (KS) tests. For post hoc analysis of ANOVAs Bonferroni’s multiple comparison test was used. To determine the presence of sub-populations within the data, for each study BLC and CA1 spine movements across all groups was combined and examined using cluster analysis. It was found that the data either clustered unbiasedly into two or three clusters. Dictating 2 or 3 clusters for each study’s cluster analysis had equal levels of good cluster quality. Visual examination of the data when grouped according to either the 2 or 3 cluster outputs revealed that 3 clusters modelled the data better than 2 clusters. Therefore, all data was analyzed using cluster analysis for 3 clusters. Cluster comparison within brain region was done using Chi squared analysis. Mann-Whitney U or Kruskal-Wallis were used to examine initial spine lengths. To examine the influence of highly motile spines on group differences, ROC was used to determine a cut off of highly motile METH spines which were then removed. Pretest training data was analyzed using Wilcoxon signed ranks tests.

## ACKOWLEDGEMENTS

We thank all members of the Miller and Rumbaugh Labs for their technical assistance and thoughtful discussions. This work was funded by grants from the National Institute Drug Abuse DA034116 (CM).

## AUTHOR CONTRIBUTIONS

EY performed the experiments, aided in the design of experiments, analyzed the data, wrote the manuscript and made the figures. HL and TK synthesized the Blebbistatin. GR trained EY, conceived of the experiments, assisted with analysis and edited the manuscript and figures. CM conceived of the experiments, assisted with analysis, wrote and edited the manuscript and edited the figures.

