## SUPPLEMENTARY INFORMATION for "Methamphetamine learning induces persistent nonmuscle myosin II-dependent spine motility in the basolateral amygdala"

**Contents:**

Supplementary Table 1

Supplementary Figures 1-5

**Table S1**

| | | Initial Spine Length | | | |
|---|---|---|---|---|---|
| | | BLA | Statistics (Mann-Whitney or Kruskal-Wallis) | CA1 | Statistics (Mann-Whitney or Kruskal-Wallis) |
| Jasp Slice Treatment | Veh | 1.56 ± 0.03 | U = 20724<br>P = 0.286 | | |
| | Jasp | 1.50 ± 0.03 | | | |
| METH CPP | Saline | 1.48 ± 0.03 | U = 28698<br>P = 0.947 | 1.21 ± 0.02 | U = 29214<br>P = 0.055 |
| | METH | 1.48 ± 0.03 | | 1.26 ± 0.02 | |
| METH CPP + Blebb Treatment | Saline/Veh | 1.66 ± 0.03 | H(3) = 7.38<br>P = 0.061 | 1.22 ± 0.02 | H(3) = 5.89<br>P = 0.117 |
| | Saline/Blebb | 1.56 ± 0.03 | | 1.24 ± 0.02 | |
| | METH/Veh | 1.64 ± 0.03 | | 1.23 ± 0.02 | |
| | METH/Blebb | 1.59 ± 0.03 | | 1.30 ± 0.02 | |

**Figure S1**

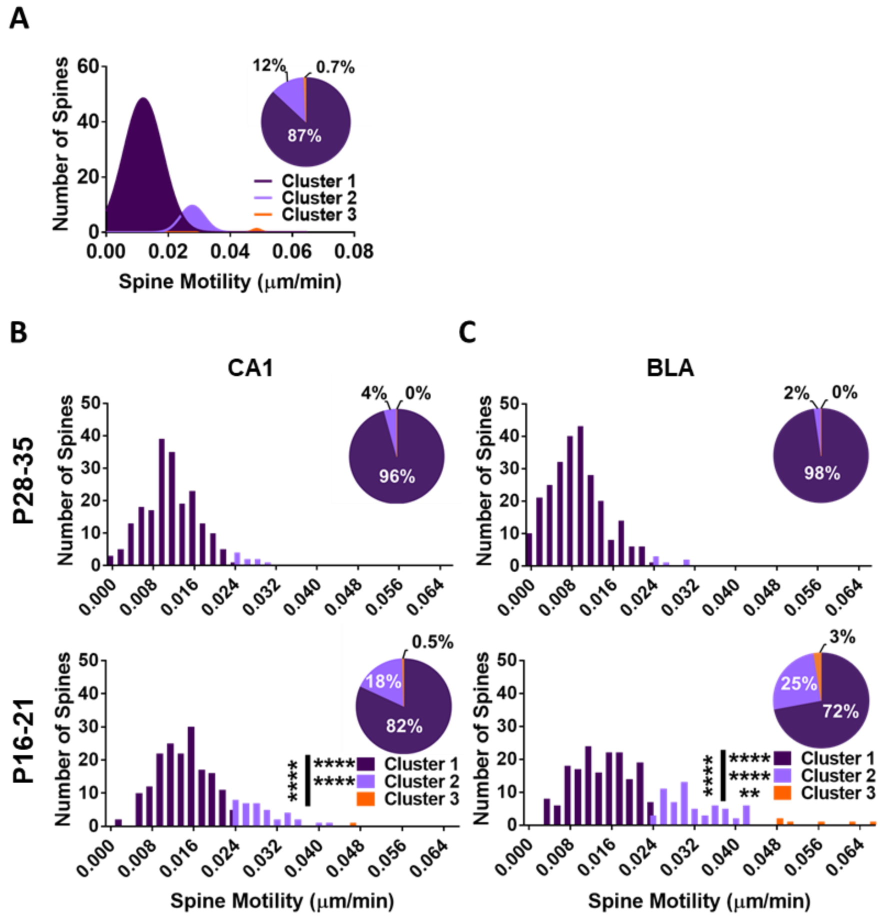


**Figure S1. Cluster analysis of CA1 and BLA age-related spine movements.** (A) All CA1 and BLA spine movements underwent cluster analysis. (C-D) Distribution of spine movements within each group across 3 clusters. Error bars represent SEM and ** *P*<0.01, *** *P*<0.001, **** *P*<0.0001.

**Figure S2**

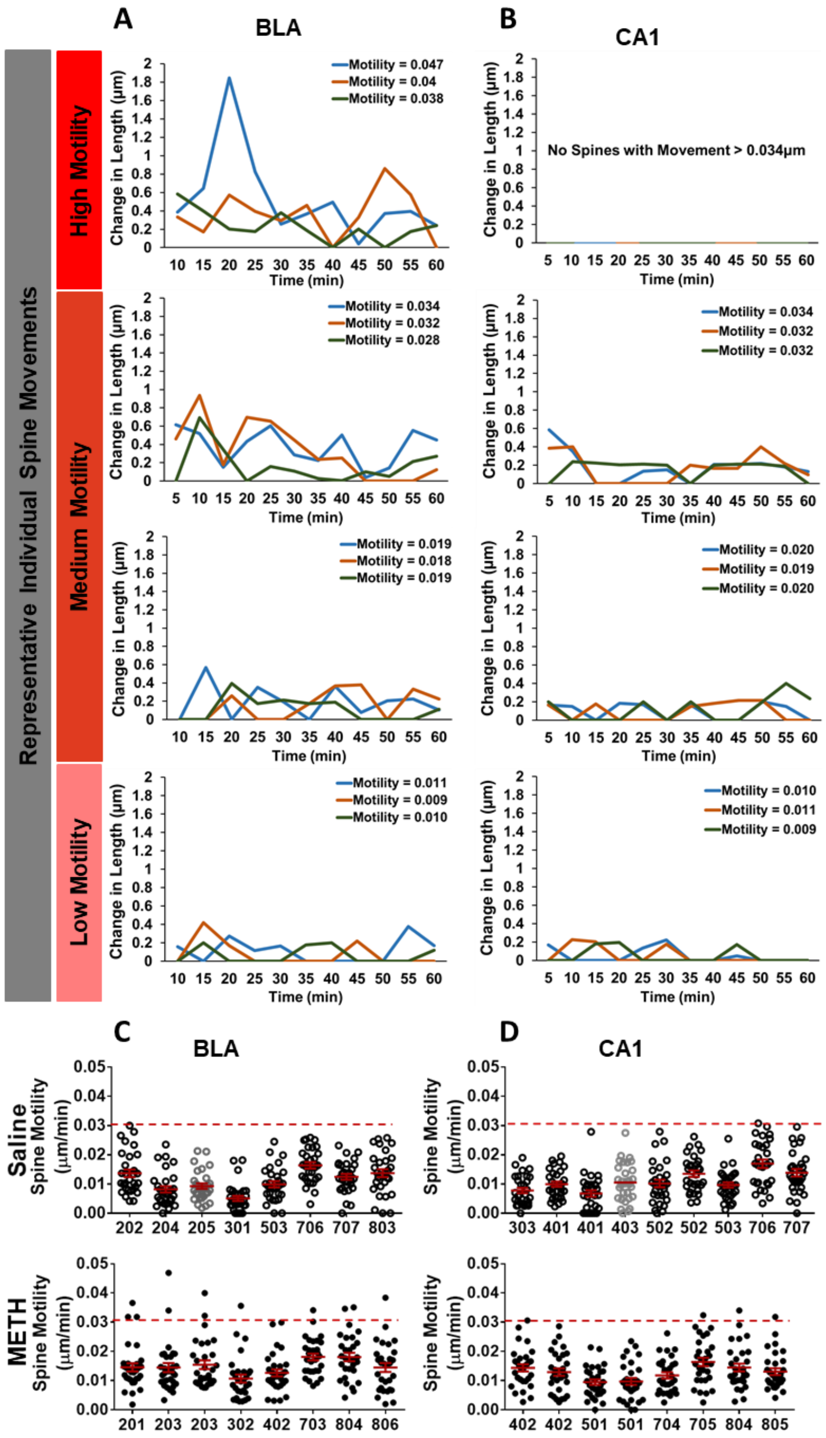


**Figure S2. BLA and CA1 spine movements following CPP training.** Representative movements of individual (A) BLA and (B) CA1 spines with different levels of motility over one hour. (C) BLA and (D) CA1 spine movements by slice and animal. Lighter colored circles denote females and darker colored circles denote males.

**Figure S3**

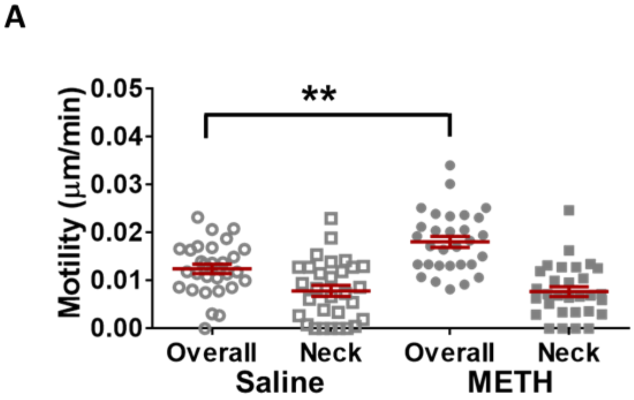


**Figure S3. Comparison of neck motility following training.** (A) Neck motility was measured in a subset of high and medium motility spines from Saline- and METH-treated animals. Error bars represent SEM and ** *P*<0.01.

**Figure S4**

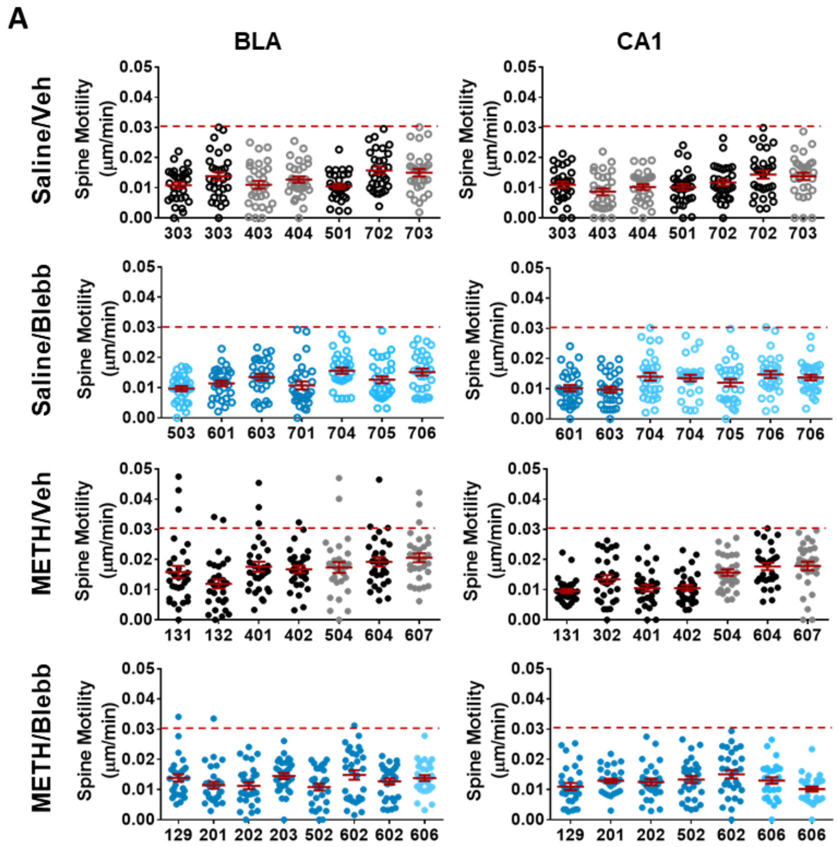


**Figure S4. CA1 and BLA spine motility across different slices and animals.** (A) 1-2 slices were used per animal and the movement of thirty spines per slice was analyzed.

### Figure S5

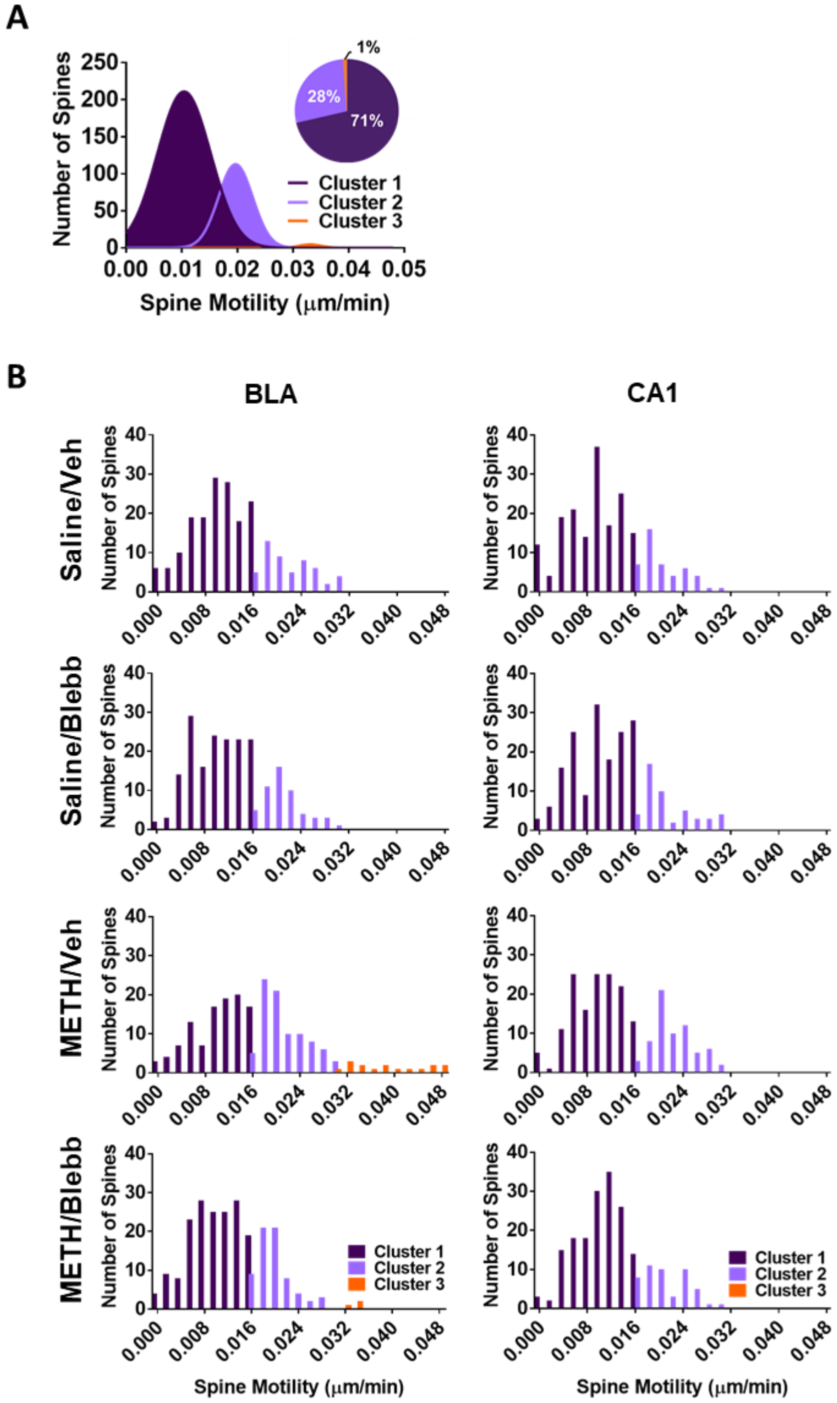


**Figure S5. Cluster analysis of CA1 and BLC spine movements following NMII inhibition.** (A) All CA1 and BLC spine movements were organized into three clusters based on cluster analysis. (B) Histograms plotting the number of spines in each cluster when the groups were graphed individually.
